# Autonomous untethered microinjectors for gastrointestinal delivery of insulin

**DOI:** 10.1101/2022.05.05.490821

**Authors:** Arijit Ghosh, Wangqu Liu, Ling Li, Gayatri Pahapale, Si Young Choi, Liyi Xu, Qi Huang, Florin M. Selaru, David H. Gracias

## Abstract

The delivery of macromolecular drugs via the gastrointestinal (GI) tract is challenging. Macromolecular drugs display low stability and poor absorption across the intestinal epithelium. While permeation-enhancing drug delivery methods can increase the bioavailability of low molecular weight drugs, the effective delivery of high molecular weight drugs across the tight epithelial cell junctions remains a formidable challenge. Here, we describe autonomous microinjectors that can efficiently penetrate the GI mucosa and deliver insulin systemically. In addition, we performed *in vitro* studies to characterize insulin release and the penetration capacity of microinjectors and measure *in vivo* release of insulin in live rats. We found that the microinjectors administered within the luminal GI tract could deliver insulin trans-mucosally to the systemic circulation at similar levels to intravenously administered insulin. Due to their small size, tunability in sizing and dosing, wafer-scale fabrication, and parallel, autonomous operation, we anticipate that these novel microinjectors could significantly advance drug delivery across the GI tract mucosa to the systemic circulation.

## Introduction

A century ago, Frederick Banting and Charles Best successfully isolated insulin from the dog pancreas and demonstrated that it could reduce blood glucose levels upon injection.^[1]^ Since then, insulin has been the mainstay in managing insulin-dependent diabetes, which currently affects more than 450 million of the global population.^[2]^ Insulin is a peptide with a molecular weight of about 5.8 kDa. Like most macromolecular drugs, it is administered by subcutaneous or intravenous injections. The injection route of insulin delivery has poor patient compliance and possible infections due to daily and repeated usage of needles, thus compromising optimal outcomes. Transdermal and oral insulin delivery routes are superior in terms of compliance and have been investigated extensively over the past several decades.^[3,4]^ Transdermal insulin delivery systems such as microneedle patches have been explored as alternatives as they are significantly less painful than hypodermic or subcutaneous needles.^[5,6]^ On applying pressure, the microneedles create minuscule disruptions in the stratum corneum, which is the main physical barrier for transporting large molecules like insulin across the skin.^[7–10]^ However, transdermal delivery using microneedle patches has proved challenging due to the difficulty of achieving therapeutic insulin levels. On the other hand, oral administration of insulin has been a long-sought-after goal because of the wide acceptance of the oral route of drug delivery.^[11–16]^ The oral route for insulin delivery is currently unavailable in clinical practice because it poses several major hurdles: (i) the passage of insulin through the mucus barrier that lines the GI epithelium, (ii) the movement of insulin across the intestinal epithelial cells held together by tight-junction proteins, and (iii) the degradation of insulin by enzymes like proteases and the acidic pH in the stomach.^[17–19]^ Over the past few decades, the use of permeation enhancers (PE) such as ethylenediaminetetraacetic acid (EDTA), glyceryl monocaprate, and sodium cholate have increased both the paracellular and transcellular transport of insulin in the GI tract^[20–22]^. However, most PEs are developed based on epithelial monolayer cultures and isolated tissue, which often results in low bioavailability in live animals and therefore has limited potential for clinical translation. ^[23,24]^

An entirely different approach to systemic drug delivery from luminal administration is to disrupt the GI epithelial tissue barrier mechanically and physically inject the drug into the subepithelial space in the vicinity of blood vessels.^[25,26]^ The method, which takes inspiration from the transdermal injection method, poses several challenges, including exerting sufficient force to penetrate epithelium inside the GI tract, unlike transdermal drug delivery, where the patch can be manually pressed against the skin. In recent years, ingestible devices demonstrating this concept include the dynamic omnidirectional adhesive microneedle system (DOAMS), the luminal unfolding microneedle injector (LUMI), and the self-orienting millimeter-scale applicator (SOMA), and the RaniPill™ capsule. These devices exploit the controlled release of energy from steel springs embedded in the device to deliver therapeutics.^[25–28]^ The relatively large size of the needles and strong forces generated by the spring in these devices pose the risk of perforating the GI tract. Also, these devices have components that are large enough to raise the possibility of GI tract obstruction, particularly in certain pathological conditions with narrowing of the GI tract, such as Inflammatory Bowel Disease.^[29]^ These devices are still in the early stage of preclinical/ clinical trials, and thus the safety and efficacy of these devices need to be evaluated.

Here, we report the development and operation of robotic, shape-changing microinjectors with an overall size of 1.5 mm when open and around 500 μm when closed, which can autonomously deliver insulin across the GI epithelium. The robotic microinjectors use thermally triggerable energy stored in prestressed thin films, effectively acting as a microspring loaded latch that can release force to enable shape change and facilitate the penetration of the tips into the epithelium (**Figure 1**a). We utilized insulin as a model macromolecular drug and incorporated insulin-loaded chitosan gel patches on the tips of the microinjectors to safely deliver insulin systemically. It is noteworthy that, unlike many larger manually assembled devices, our microinjectors can be fabricated using wafer-scale processes, like those used in the semiconductor industry, and are scalable across sizes. Moreover, the microinjectors are small enough to be used in large numbers without causing any GI blockage or visible trauma in the animals. Our proof-of-concept studies in rodents show, for the first time, that shape-changing miniaturized injectors administered enterally can safely and systemically deliver a therapeutic dose of insulin, similar to that from intravenous injection.

**Figure 1.**
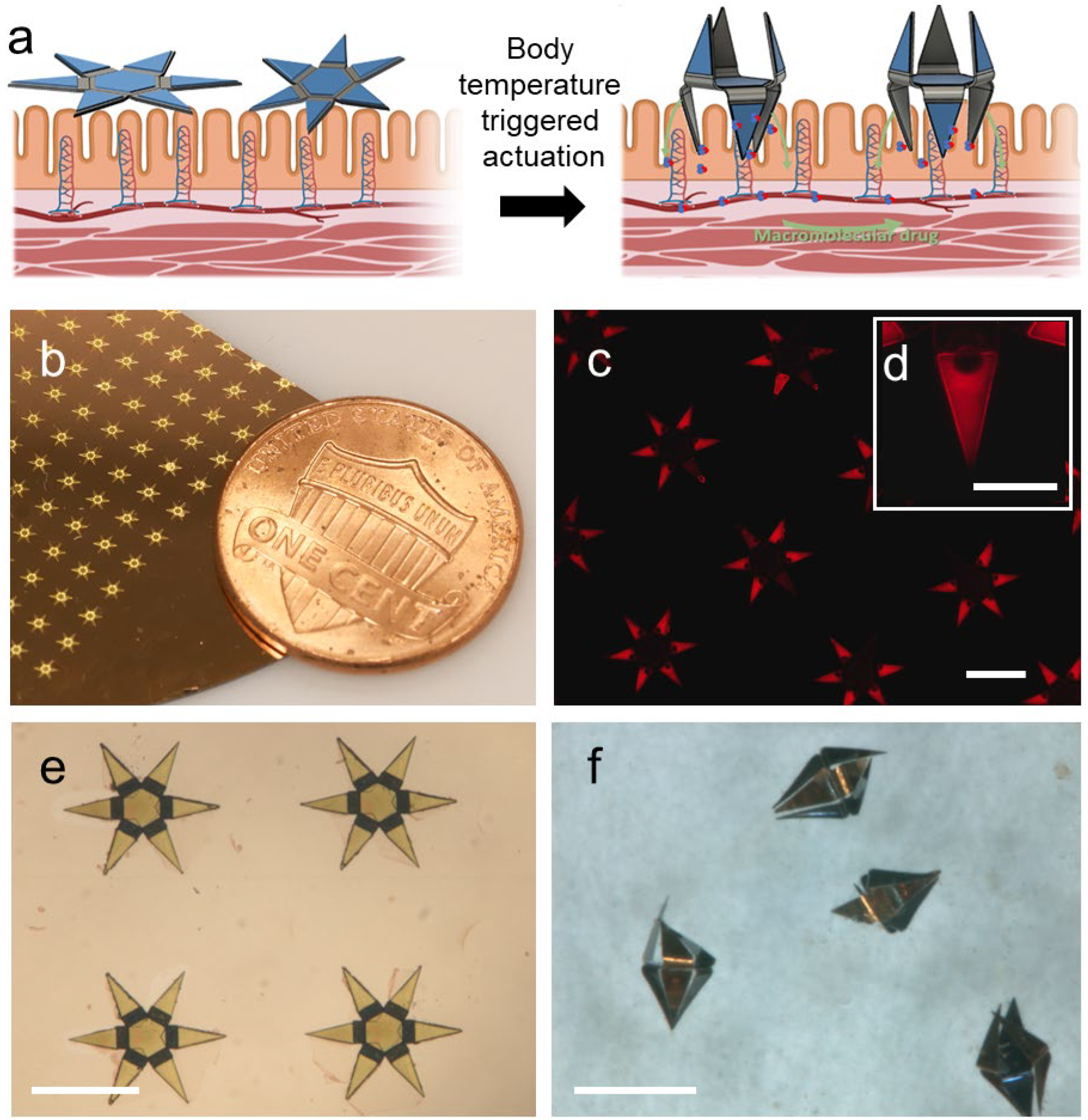
Design, fabrication, and operation of autonomous microinjectors. (a) Conceptual illustration showing autonomous actuation of two microinjectors with their injection tips penetrating the mucosa as the injectors equilibrate to physiological temperature, while the macromolecular drugs are transported across the mucosal epithelium. (b) Photo of microinjector arrays fabricated on a silicon wafer near a penny, illustrating the small size of the microinjectors and parallel wafer-scale fabrication. (c) Fluorescence image of an array of as-fabricated microinjectors loaded with fluorescent rhodamine within the chitosan to aid visualization of the gel patch on the injection tips. The scale bar is 1 mm. (d) The inset shows a zoomed fluorescence image of a single injection tip loaded with fluorescent rhodamine within the chitosan. The scale bar is 200 μm. (e-f) Optical microscopy images of microinjectors; (e) as fabricated on a silicon wafer. And (f) bidirectionally folded post actuation in response to a physiological temperature. The scale bars are 1 mm.

## Results and Discussion

We designed the robotic microinjectors using origami design principles ^[30–33]^ and the designs consist of hinge and tip segments. The hinge segments of the microinjector generate the injection force necessary for the tips to penetrate the tissue. The injection force is produced by the thermally triggered release of intrinsic differential stress in thin-film multilayers of chromium (Cr) and gold (Au) at the hinge. Each microinjector is fabricated as a 1.5 mm tip-to-tip 2D multilayer device, which can self-fold to form a 500 μm 3D device. The microinjectors are equipped with six injection tips, 450 μm in length, which are coated with a mucoadhesive chitosan gel loaded with insulin (Figure 1). Notably, we incorporated a bidirectional foldable design of the microinjectors that allows the injection tips to deliver the drug in any direction, irrespective of the orientation in which they land on the GI epithelium.

Each microinjector is a multilayer thin-film structure consisting of five layers (Figure S1a). We used computer-aided drawing (CAD) to design photomasks for patterning individual layers during the fabrication process and can accommodate 483 microinjectors on a 3-inch diameter silicon wafer. This number can be scaled up easily to several thousand per wafer if the fabrication is carried out on a 12” diameter wafer, which is routinely used in semiconductor foundries, thus further bringing down the fabrication cost. Figure 1b shows fabricated microinjectors aside a penny indicating their small size. We achieved folding bidirectionality of the microinjection tips by engineering the design of the multilayer thin film stacks such that hinges could bend in opposite directions (Figure S1b). We fabricated microinjectors with this design by depositing two different thin-film multilayer assemblies: (i) a two-layer assembly of Cr/Au and (ii) a four-layer assembly of Cr/Au/Cr/Au.

We estimated the relative thickness of the layers in these designs using a theoretical model which optimizes fold angle based on the thickness, differential stress, modulus, and Poisson ratios (Details in Note S1). On top of the differentially stressed multilayers, we electrodeposited thick nickel (Ni)/Au rigid panels on the center and tip segments. The microinjector regions with such rigid panels do not bend, while the thinner hinges bend and fold when actuated. We spatially patterned insulin-loaded chitosan gel patches on the microinjector tips using a combination of photolithography and electrodeposition, as described in a previous study (Figure 1c, d). ^[33]^ Finally, we patterned a paraffin wax trigger layer atop the hinges of the microinjectors; this layer softens at the physiological temperature of the GI tract and acts as the thermal trigger to induce the bending of the hinges and folding of the microinjectors. A schematic of the entire fabrication process flow is shown in Figure S1c. After fabrication, we released the microinjectors from the silicon wafer and stored them at room temperature (approximately 23 °C) or in a refrigerator (approximately 4 °C). When administered from the cold state, we observed that the injection tips activate within a few minutes once the microinjectors equilibrate with the physiological temperature of the GI tract (Figure 1f, Movie SM1).

The microinjection tips with insulin-loaded chitosan gel layers of the microinjectors were approximately 5 μm in thickness (**Figure 2**a-b). This thin size of microinjector tips is necessary to generate a high pressure to penetrate tissue at the injection site. We theoretically estimated the maximum pressure exerted by the microinjection tips to be 0.4 - 0.5 MPa for the tips with the drug-loaded chitosan layer and 0.5 - 0.6 MPa for the tips without chitosan gel using the Hertz contact mechanics model (Note S3). Note that although the dimensions and design are different, the pressure exerted by the microinjector tips is consistent with previously described theragrippers. ^[33]^

**Figure 2:**
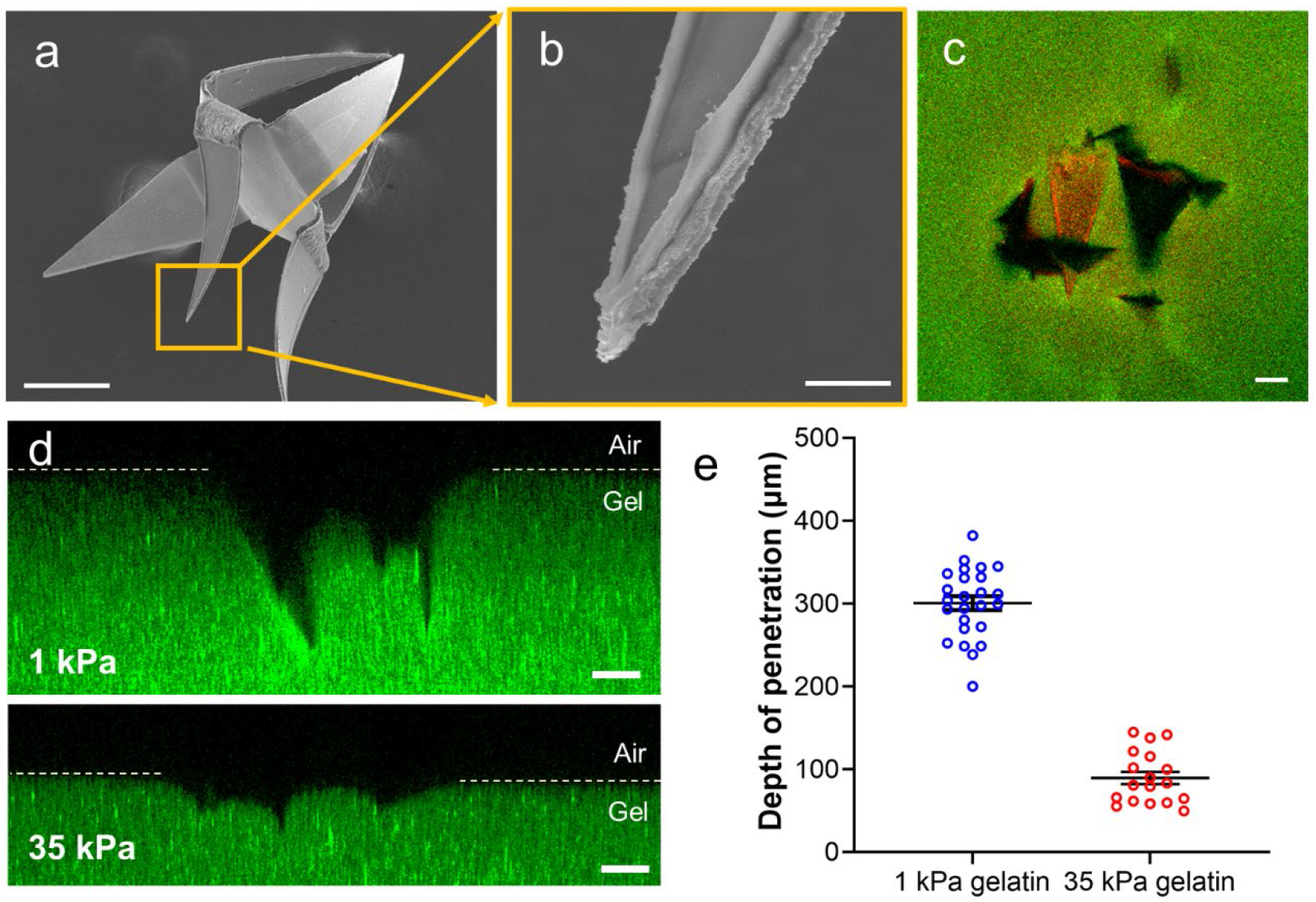
Evaluation of the penetration of the microinjector tip into tissue-mimicking gelatin. (a) Scanning electron microscope (SEM) image of a bidirectional microinjector after actuation, illustrating that the injector arms fold in opposite directions. The scale bar is 200 μm. (b) Magnified SEM image of a microinjector showing the insulin-loaded chitosan patch on the tip. The scale bar is 10 μm. (c) Confocal image showing the top view of the microinjector (the chitosan gel on microtips are loaded with rhodamine dye for visualization) penetrating a 1 kPa gelatin hydrogel (loaded with 200 nm diameter fluorescent polystyrene beads for visualization). The scale bar is 100 μm. (d) Cross-sectional confocal fluorescence microscopy image (side view) of the microinjector tips penetrating a 1 kPa (top) and a 35 kPa (bottom) gelatin hydrogel. The images show that the microinjection tips penetrate significantly deeper into the soft 1 kPa gelatin. The scale bars are 100 μm. (e) Plot depicting the depth of penetration of the microinjector tips into the gelatin hydrogels of two different stiffnesses, 1 kPa (blue) and 35 kPa (red). Data presented were measured for each of the three penetrated microtips from at least 4 samples, and the plot shows the mean and standard error of the mean.

We evaluated the injection performance of the microinjectors on gelatin hydrogels having a stiffness of 1 kPa, which is close to the stiffness of the colonic mucosa (∼0.7 kPa) (Figure 2c). ^[34]^ The preparation of thermally stable gelatin has been described in the Materials and Methods section. ^[35]^ For ease of visualization, we used rhodamine dye as a model drug. We actuated the microinjectors by placing them in an oven set at 40 °C for 15 to 20 minutes (Figure S3) to simulate the physiological temperature. We observed that the microinjector tips penetrated approximately 300 μm into the 1kPa gel. (Figure 2e). We also conducted a similar experiment with a significantly stiffer gelatin hydrogel (35 kPa) to estimate the microinjection tip penetration depth in a stiff tissue environment (Figure 2d). We found that the microinjector tips could only penetrate up to 100 μm into the 35 kPa gel while it could penetrate up to 3 times the depth into 1kPa gelatin biomimetic hydrogel. (Figure 2e).

The performance of the microinjectors was then evaluated on *ex vivo* rat colon tissue. We placed the microinjectors on top of freshly excised rat colon tissue which was incubated in a Petri dish covered with saline (**Figure 3**a-b). We heated the tissue and injectors assembly in an oven to 40 °C (Figure 3c) and evaluated the tissue penetration abilities of the microinjectors using scanning electron microscopy (SEM) and micro-computed tomography (μ-CT). As shown in Figure 3d-g, the microinjectors penetrated approximately 250 μm into the rat colon mucosa, which is consistent with the gel penetration experiment results. We also conducted similar experiments on a freshly excised pig stomach and colon, and the results are shown in Figure S4.

**Figure 3:**
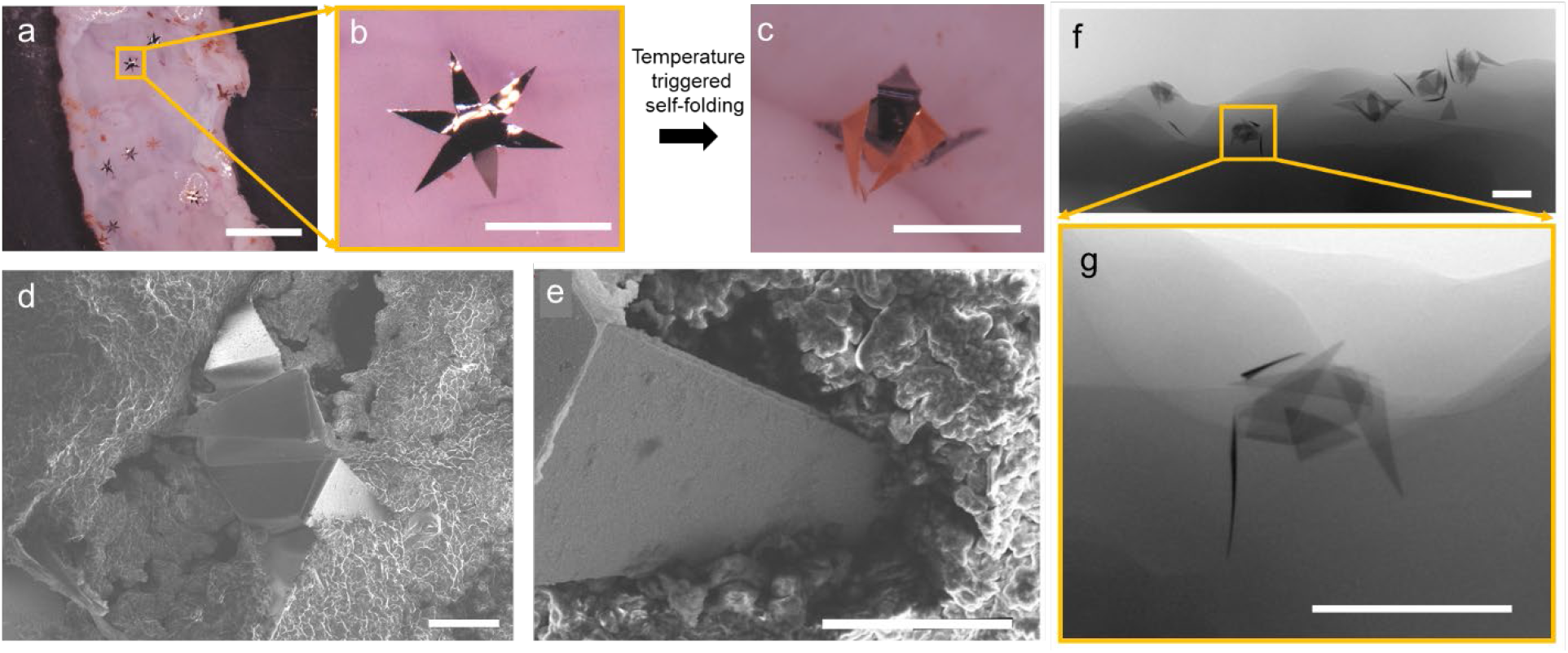
Autonomous operation of the microinjectors on *ex vivo* rat colon. (a) Optical image showing microinjectors before actuation on freshly excised rat colon tissue *ex vivo*. The scale bar is 5 mm. (b-c) Zoomed images of a microinjector, (b) before, and (c) after autonomous actuation at physiological temperature. The scale bars are 1 mm. (d) SEM image of a microinjector attached to the rat colon tissue. The scale bar is 200 μm. (e) SEM image showing the penetration of an injection tip into the colon tissue. The scale bar is 10 μm. (f) µ-CT image of an *ex vivo* rat colon with microinjectors attached to it. The scale bar is 1 mm. (g) Zoomed in µ-CT image of the microinjector marked in panel f, showing the depth of penetration into the tissue. The scale bar is 1 mm.

After demonstrating that the microinjectors could penetrate deep into the colon tissue *ex vivo*, we verified the tissue-penetrating capability of the microinjectors *in vivo* in live rats. For these experiments, we used 280 - 350 g male Wistar rats, which have a typical colon diameter of approximately 8 mm. We administered approximately 200 microinjectors through the rectum into the colon of these rats (Figure 4a) using a pneumatic microfluidic controller, which could eject a bolus of the microinjectors in saline using controlled pressure. We used a pressure of 14 - 16 psi with a medical-grade polycarbonate tubing with an internal diameter of 2.5 mm, to drive the bolus of microinjectors into the rat colons. We observed no adverse effect on the health of the animals during and even 48 hours after the deployment of the microinjectors. Figure 4b-c show μ-CT images of microinjectors still presented and attached to the colon of rats 48 hours after their intrarectal administration.

**Figure 4:**
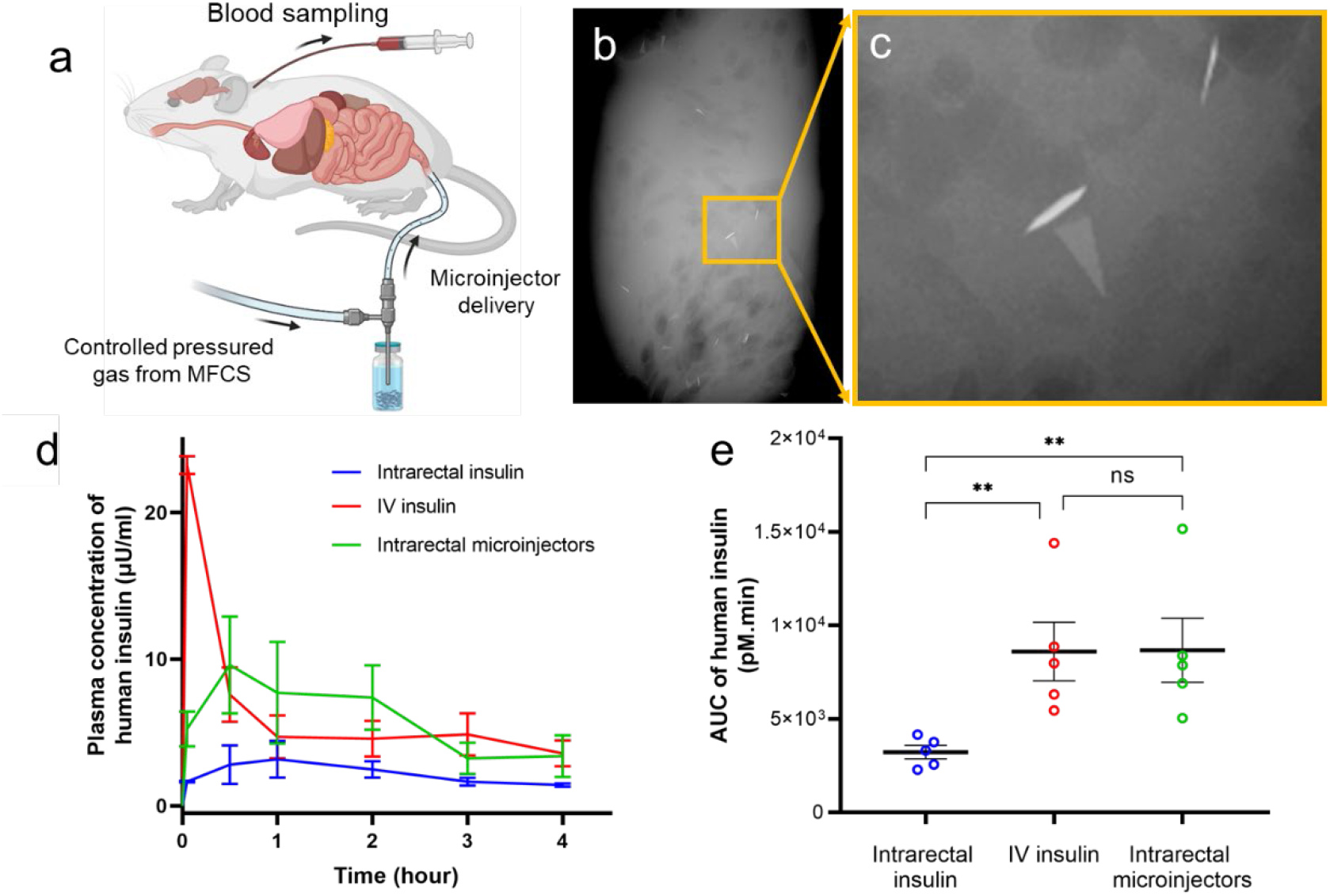
Enteral delivery of human insulin using microinjectors in live rats. (a) Schematic of the *in vivo* experiments, which involve the intrarectal administration of human insulin formulated microinjectors. The microinjectors were delivered at a controlled air pressure using a microfluidic controller. (b-c) µ-CT image of the excised colon, 48 hours post rectal administration in rats, showing the presence of microinjectors attached to the colon (d) Pharmacokinetic (PK) profile showing the concentration (mean and standard error of the mean) of human insulin measured in rat plasma after administration over four hours. In these studies, the insulin was injected intravenously (red, positive control), intrarectally using microinjectors (green), and intrarectally (blue, negative control) using the microfluidic controller. Each experimental arm was conducted for five different male rats. (e) Plot showing the comparison of the area under the PK curves in panel (d). The plot shows the significant advantage of using microinjectors to achieve a similar level of insulin exposure to IV insulin. The plot shows the mean and standard error of the mean (N=5).

The microinjectors’ function as drug delivery vehicles was tested by using insulin as a model macromolecular drug in a live rat model, as insulin has been a commonly used model drug in the past. ^[21]^ We loaded insulin into the microinjector tips by soaking them in a concentrated insulin solution (see Materials and Methods).

First, we studied the *in vitro* release profile of insulin from the microinjectors over 4 hours. We placed the 200 insulin-loaded microinjectors in saline at 37 °C and measured the released insulin in the solution over time. Based on these measurements, we estimate that each microinjector has the capacity to accommodate around 300 μIU of human insulin. Moreover, these measurements indicated the microinjector could steadily release insulin *in vitro* (Figure S5).

We then conducted *in vivo* insulin delivery experiments, in which 60 mIU of human insulin was administered per animal. Each experimental arm consisted of 5 rats and is described as follows: In the first arm, we delivered an intrarectal dose of 60 mIU of human insulin in 1 mL of saline (negative control). We delivered an intrarectal dose of 60 mIU of human insulin in the second arm, formulated with 200 microinjectors (experimental arm). We carried out the rectal administration for both these groups using a pneumatic microfluidic controller. We administered an intravenous (IV) dose of 60 mIU of human insulin in the third group through a jugular vein catheter (positive control). We then drew blood from all the animals at t = 5 min, 30 min, 1 hr, 2 hr, 3 hr, and 4 hr post administration of the insulin. We used a commercial enzyme-linked immunosorbent assay (ELISA) to determine human insulin concentration at various time points over the 4-hour time window. The pharmacokinetic (PK) profile of human insulin in rat plasma over 4 hours after administration is shown in Figure 4d. We measured peak plasma human insulin concentration administered by the microinjectors in the plasma of rats to be 9.6 μIU/ml at the 30 min time point. In contrast, the rats with intrarectally administered insulin solution without microinjectors (negative control arm) had only a minimal amount of human insulin in their plasma. Furthermore, we measured the total exposure of insulin by calculating the area under the PK curves. The results are plotted in Figure 4e, where we see that the microinjectors provide similar total exposure of insulin in the bloodstream compared to the IV-dosed animals, although with a different PK profile. As expected, animals treated with IV insulin had a sharp increase in plasma level of insulin that then dropped precipitously. Animals treated with insulin-loaded microinjectors showed a slower increase but more sustained release, which is likely a function of absorption from the submucosal space into the systemic circulation.

We examined the insulin delivery efficiency of our microinjectors and compared that to other GI tract administered insulin delivery mechanisms. We made the comparison in terms of the maximum insulin plasma concentration as well as insulin dosage per body surface area (BSA) of the animals (details in Note S5). The microinjectors show a significantly higher insulin bioavailability in the rat model over other GI tract-administered insulin vehicles. Specifically, we measured the highest human insulin plasma concentration of 65.3 pM in rats that received insulin-loaded microinjectors, with 0.063 mg/m^2^ BSA initial dosage. We divided the highest plasma insulin concentration by the initial dosage to get a normalized insulin delivery coefficient of 1036.5 pM/mg·m^-2^ BSA for microinjectors. In comparison, the delivery coefficients in previously reported insulin delivery vehicles (studied in a rat GI tract) are as follows: (i) alginate/chitosan nanoparticles - 9.2 pM/mg·m^-2^ BSA, (ii) HEMA nanogels - 9.5 pM/mg·m^-2^ BSA, and (iii) hydrogel patches - 54.7 pM/mg·m^-2. [36–39]^ We attribute the high insulin delivery efficiency to the fact that the microinjector tips penetrated the GI mucosa, which greatly enhanced the macromolecular drug diffusion. Furthermore, we compared the insulin delivery efficiency of our microinjector with other GI tract based insulin injection devices studied in the pig model. For example, the SOMA device, which operates in the stomach, has a delivery coefficient of 111.1 pM/mg·m^-2^. The LUMI device that works in the small intestine shows a delivery coefficient of 81.8 pM/mg·m^-2^.^[25,26]^ Although the LUMI and SOMA devices show a better delivery efficiency as compared to the nanoparticles, hydrogels and patches discussed previously, the microinjectors reported in this study outperform them by at least an order of magnitude. We attribute this outperformance to the the large number of up to 600 different microinjection sites in our study as compared to one^[25]^ or tens of^[26]^ injection sites created by other larger GI injection devices. The ability of the microinjectors to perform autonomous injections in small conduits like a rat GI tract also suggests the possibility of accessing narrower conduits than the GI tract to perform localized drug delivery.

In summary, oral administration of macromolecular drugs such as insulin for systemic delivery would dramatically improve patient outcomes and reduce costs by increasing compliance and decreasing complications and hospitalizations. However, enhancing the diffusion of these drug molecules across the GI epithelium is challenging. Here, we have introduced a new concept of using miniaturized microinjectors, which are small enough to be safely ingested and can significantly enhance the transportation of macromolecule drugs like insulin across the GI tract. As the self-injecting device is independent of the encapsulated active molecule, the microinjector platform can be potentially formulated to deliver even fragile drugs such as peptides, antibodies, and RNA, which rarely have oral formulation. However, it remains to be seen how the design of the microinjectors can be scaled up for efficient delivery in larger animal models^[25],^ which is essential for successful translation to the clinic. Though our method was found to be safe in general, we envision that the use of transient and biodegradable materials to fabricate the microinjectors will further enhance the biocompatibility and safety ofthe proposed method of delivery. ^[40-41]^

## Materials and Methods

### Fabrication of the microinjectors

We fabricated the microinjectors using planar microfabrication techniques on silicon wafers (Figure S1a). First, we deposited a sacrificial layer of copper (Cu, 300 nm) with underlying chromium (Cr) adhesion layer (20 nm) using thermal evaporation. The microinjector fabrication that followed on the top of the Cu layer consists of a combination of photolithography, thermal evaporation of Cr and gold (Au), electrodeposition of nickel (Ni), Au, and chitosan, and spin coating steps for the photoresist and wax. We fabricated microinjectors that actuate in one direction (unidirectional) or two directions (bidirectional) by incorporating either one or two differentially stressed Cr/Au layers. For the bidirectional microinjector, each of the stress layer assemblies consists of three alternatively arranged microinjector tips. The details of the microinjectors design are described in Supporting Note S1. We fabricated the tips by creating a photolithographically defined pattern of six (for unidirectional) or three (for bidirectional) injection tips using S1813 photoresist (MicroChem Corp.) on copper. We patterned the first stress layer assembly by evaporating 60 nm Cr/100 nm Au and lift-off metallization. We then patterned the three alternate injection tips using a second photolithography step using the S1813 photoresist. This assembly consists of 15 nm Cr / 100 nm Au / 75 nm Cr /10 nm Au. After the stress layer deposition, we did a photolithography step using the SPR™ 220 photoresist (Megaposit™, Kayaku). We created the rigid panels on the differentially stressed multilayer assemblies by electroplating 3 μm of Ni and 0.3 μm of Au using commercial nickel sulfamate and gold sulfite solutions (Technic). ^[42]^ After photoresist stripping, we patterned a photoresist mold layer using photolithography on the microinjection tips. Using electrodeposition, we filled the mold using chitosan (medium molecular weight, Sigma-Aldrich). We dissolved the photoresist mold (within 24 hours) using acetone. We then patterned the paraffin wax (melting point 53 to 58 °C, Sigma-Aldrich) trigger layers on the hinges of the microinjectors using another step of photopatterning SPR™ 220 photoresist on the hinges of the microinjectors. To ensure proper coverage of paraffin wax on the hinges of the microinjectors, we optimized the volume of wax dropped on the wafer and the spin coating conditions. Further details of the fabrication optimization studies are in Note S2, Figure S2, and Table S2. After the deposition of paraffin wax, we allowed the wax to sit for at least two hours, and then we dissolved the photoresist. We then released the microinjectors from the wafer by dissolving the Cu sacrificial layer in a commercial basic cupric chloride solution (copper etchant BTP, Transene), which preserves the chitosan patch on the injection tips and the paraffin wax on the hinges. We thoroughly rinsed the injectors in DI water to remove any residual etchant.

### Gelatin penetration experiments

We prepared microbial transglutaminase (mTG) crosslinked gelatin hydrogels for the evaluation of the microinjector penetration as described previously. ^[35]^ Briefly, we dissolved 12.5% by weight gelatin powder (Type A, porcine skin, Sigma Aldrich) in PBS and sterile filtered it through a 0.2 μm polystyrene membrane filter. We mixed the gelatin solution with 1 mL of sterile-filtered 10 U/g-gelatin of mTG (Ajinomoto) prepared in PBS. We used Bloom 90-110 (low molecular weight, 20-25 kDa) and Bloom 300 (high molecular weight, 50-100 kDa) gelatin to prepare the 1 kPa (soft) and 35 kPa (stiff) hydrogel, respectively. We mixed 200 nm green, fluorescent polystyrene beads (Polysciences) with 350 μL of the gelatin-mTG solution to aid in visualization and added them to dishes that had a 20 mm glass bottom (MatTek Corporation). We allowed the gel to crosslink for 8 hours at 37 °C. After the crosslinking reaction was completed, we heated the gelatin gels in PBS to 60 °C for 30 mins to deactivate the mTG and stored them in PBS at 37 °C until subsequent use in the experiments.

To estimate the penetration of injector tips, we placed the microinjectors on the gelatin hydrogel surface under PBS and placed the gelatin in an oven set at 40 °C. We observed that the wax trigger layer softened at this temperature and induced the folding of the injectors on the gelatin hydrogel surface. We imaged the microinjectors using a 10x objective on an A1 confocal microscope (Nikon) and estimated the injector tip penetration depth from the confocal z-stack using ImageJ. We created an orthogonal projection (side/xz view) of the confocal z-stack for each injector tip and measured the penetration depth from the hydrogel surface.

### Ex vivo tissue penetration experiments and imaging

We used freshly excised rat colon and pig colon, stomach, and esophagus to carry out our *ex vivo* tissue penetration experiments with the microinjectors. We euthanized 300 g male Wistar rats (Charles River, MA) and removed the colon. We cleaned the colon and laid it flat to face up on the luminal side. We incubated the flat colon sections under saline and dropped microinjectors on the top of the tissue samples. We then placed the microinjector/tissue assembly in an oven at 40 °C for 15 to 20 minutes. The microinjectors actuate at the increased temperature and penetrate their injection tips into the tissue. We performed similar experiments with pig colon, stomach, and esophagus. We procured the pig organs from freshly sacrificed animals from a butcher shop (Wagner Meats LLC, Mount Airy).

We characterized the penetration of microinjectors on *ex vivo* tissues using optical microscopy, scanning electron microscopy (SEM, JEOL), and microcomputed tomography imaging (μ-CT, RX Solutions). To image the tissue samples with microinjectors, we prepared the samples as follows: For optical microscopy and μ-CT, we used the fresh tissue, without any drying and fixing, to preserve the surface characteristics of the mucosa as much as possible. For SEM imaging, we collected the tissue samples with microinjectors tips penetrated into it. Then we used sodium cacodylate buffer to wash the tissue samples and fixed them in glutaraldehyde for an hour. We then washed the tissue in sodium cacodylate buffer and post-fixed in osmium tetroxide for one hour on ice and in the dark. Afterward, we rinsed the tissue samples in DI water and performed the tissue dehydration using a graded series of cold ethanol washes (50%, 70%, 90%, and 100%) for 15 minutes each. We then successively soaked the samples at room temperature in the following solutions: anhydrous ethanol for 20 minutes two times, a mixture of 50% hexamethyldisilazane (HMDS), and 50% anhydrous ethanol for 30 minutes, and finally pure HMDS for 30 minutes. We air-dried the samples before putting them in the SEM vacuum chamber.

### In vitro measurements of insulin release from microinjectors

To prepare the insulin-loaded microinjectors, we released around 100 microinjectors from the silicon wafer by dissolving the Cu sacrificial layer and rinsing at least six times to remove the Cu etchant and obtain a clear solution of microinjectors in DI water. We then replaced the DI water with 5 mg/mL of human insulin saline solution, in which we soaked the microinjectors for 36 hours at room temperature (around 23 °C). After that, we washed the microinjectors with DI water at least six times to remove the excess insulin.

We conducted the *in vitro* human insulin release experiments by immersing around 100 microinjectors in 10 mL of saline at around 37 °C. At each desired time point (Figure S5), we withdrew 100 μL from the solution and replaced it with 100 μL of fresh saline to maintain a proper sink condition. We then measured the concentrations of human insulin in the samples at various time points using a commercial ELISA kit (ALPCO, 80-INSHU-E01.1) and a spectrophotometer (Molecular Devices, SpectraMax i3). We repeated the experiment thrice and plotted the cumulative concentrations in Figure S5.

### In vivo experiments to deliver human insulin using microinjectors

We prepared the human insulin-loaded microinjectors for *in vivo* animal experiments using the same method described in the *in vitro* experiment section above. We used 200 (within 2%) microinjectors for each animal. We performed the *in vivo* experiments on male Wistar rats with a jugular vein catheter weighing approximately 300 g (Charles River, MA). The experiments followed the Johns Hopkins University Animal Care and Use Committee protocol number RA19M207. We fasted the rats for one day before the experiments for an empty colon. We mildly anesthetized the rats using isoflurane and oxygen while intrarectally administering the microinjectors. We stored the human insulin-loaded microinjectors in 2 mL vials. To deliver the microinjectors, we attached a medical-grade polytetrafluoroethylene (PTFE) tube with a 2.5-mm-inner diameter (Zeus Inc.) to a computer-controlled pneumatic delivery system (Fluigent, MFCS-100 1C). We inserted it 3 to 4 cm inside the colon of the animal. We ejected the microinjectors with a small amount of DI water at 14-16 psi pressure. We returned the rats to the cage after microinjector administration.

We drew a 100 μL blood sample via the jugular vein cannula for bioanalysis at the predefined time points. We mixed the blood samples with 20 IU heparin and then centrifuged them at 3000 rcf for 10 min to separate the plasma. We stored the resulting plasma at −80 °C until the insulin assay measurements.

### Assay for the detection of human insulin in rats

We used an enzyme-linked immunosorbent assay (ELISA) to conduct the insulin concentration measurements following the manufacturer-directed assay procedure. Briefly, to determine insulin concentration in rat plasma, we used an ultrasensitive human insulin-specific ELISA kit (ALPCO, 80-INSHUU-E01.1), having a sensitivity of 0.135 μIU/mL and a dynamic range of 0.15-20 µIU/mL, which is insensitive to rat intrinsic insulin. To determine insulin concentrations in saline in our *in vitro* release experiments, we used an ELISA kit (ALPCO, 80-INSHU-E01.1) with a sensitivity of 0.399 µIU/mL and a dynamic range of 3.0-200 µIU/mL. We used a microplate shaker (800 rpm) to react to the rat plasma samples and the detection antibody in a 96 well-plate, which is precoated with a monoclonal antibody specific to human insulin. We duplicated each plasma sample during the measurement for each time point and each animal. We used rat plasma collected before the insulin administration as the control and used the manufacturer-provided standard solutions during the analysis. After the reaction with the antibodies, we washed the wells thoroughly with a buffer. We then used colorimetric detection to measure the absorbance of each well at 450 nm wavelength with a spectrophotometer (Molecular Devices, SpectraMax i3). We compared the absorbances with a previously obtained standard curve of human insulin (Figure S6), acquired using the manufacturer-provided standard solutions. We plotted the extracted insulin concentrations as a function of the time of sample collection (Figure 4d) and the area under the curve (Figure 4e) for the different experiments. Please see Note S4 for further details about the insulin dose determination and assay validation procedure (Figure S7).

## Supporting Information

Supporting Information is available

## Acknowledgments

We acknowledge support from the National Institute of Biomedical Imaging and Bioengineering of the National Institutes of Health under Award Number R01EB017742. The content is solely the authors’ responsibility and does not necessarily represent the official views of the National Institutes of Health.

## Author Contributions

AG, DHG, and FMS conceptualized the study and designed the experiments; DHG and FMS supervised the study; AG, LX, and SC performed the microfabrication and *in vitro* device characterization; GP carried out the gel imaging studies; AG, WL, LL, and LX performed the animal experiments; AG, LX, and QH did the SEM and µ-CT imaging of tissues; AG did the bioanalysis, and pharmacokinetic data analysis; AG and WL made the illustrations; AG, WL, FMS and DHG wrote the manuscript with input from all authors.

