## Supplementary Information for "Autonomous untethered microinjectors for gastrointestinal delivery of insulin"

Dr. A. Ghosh, W. Liu, G. Pahapale, S.Y. Choi, L. Xu, Q. Huang and Prof. D.H. Gracias
Chemical and Biomolecular Engineering, Johns Hopkins University, Baltimore, MD 21218, USA

Dr. L. Li, Dr. F.M. Selaru
Gastroenterology and Hepatology, Department of Medicine, Johns Hopkins University, Baltimore, MD 21287, USA

Prof. D.H. Gracias
Materials Science and Engineering, Johns Hopkins University, Baltimore, MD 21218, USA

Prof. D.H. Gracias
Department of Chemistry, Materials Science and Engineering, Laboratory for Computational Sensing and Robotics (LCSR), Johns Hopkins University, Baltimore, MD 21218, USA

Prof. D.H. Gracias, Sidney Kimmel Comprehensive Cancer Center (SKCCC), Department of Oncology, Johns Hopkins University School of Medicine, Baltimore, MD, USA

### Supporting Notes

#### Supporting Note S1: Design of the microinjectors

We designed bidirectional microinjectors with tips that fold in opposite directions to ensure that they can penetrate the gastrointestinal (GI) mucosa, irrespective of the orientation of the injectors. We used two separate and alternating stress layer assemblies in the fabrication process to drive the microinjection tips in opposing directions (Figure S1). The first assembly comprises three injection tips connected by a Y-shaped center. The second assembly comprises three other injection tips connected by a hexagonal-shaped center. Compared to designs with two hexagonal-shaped centers, the design that used a combination of a Y-shaped and hexagonal-shaped center significantly reduced microinjector breakage.

We used a multilayer thin-film model by Nikishkov [1] to estimate the desired thickness values of the chromium (Cr) and gold (Au) layers in the fabrication of the stress layer assemblies. The relevant equations can be found in reference. [2] In the calculation, we assumed the stress values of Cr to be 1 GPa and that of Au to be zero.[2] We calculated the bending curvature and, by extension the folding angle of a specific length of the multilayered hinge, which drives the actuation of the microinjection tips. It may be noted that, while in principle, stress layer assemblies having Cr 60 nm/Au 100 nm and Au 100 nm/Cr 60 nm should produce opposing foldable injection tips, we need to add an adhesion-promoting Cr and an Au seed layer for electroplating which required us to incorporate a two-layer and a four-layer assembly to fabricate the microinjectors. We used 15 nm Cr as the first layer to improve the adhesion of the assembly to the Cu sacrificial layer and a 10 nm Au thin film as a seed layer to enhance adhesion and prevent delamination of electroplated films of Ni from the evaporated Cr films. The first stress multilayer layer assembly thickness was kept constant as Cr 60 nm/Au 100 nm, and the thickness for the second stress multilayer assembly is shown in Table S1.

**Table S1. Thickness values used in the second stress layer for the fabrication of our microinjectors from the bottom to the top.**

| Trials \ Thickness | Cr/nm | Au/nm | Cr/nm | Au/nm | Fabrication yield | Predicted folding angle/° |
|---|---|---|---|---|---|---|
| **1** | 10 | 100 | 70 | 5 | 1.70% | -247.6 |
| **2** | 15 | 100 | 75 | 5 | 26.10% | -201.1 |
| **3** | 15 | 100 | 75 | 10 | 99.20% | -176.7 |

**Supporting Note S2: Optimization of the thermoresponsive trigger layer deposition**

We spin-coated a paraffin wax layer on top of the hinges as the thermoresponsive trigger. The paraffin wax used remains stiff at low temperature and softens at the physiological temperature allowing the injection tip hinges to actuate. However, improper patterning coverage of the paraffin on the hinges (Figure S2) results in premature actuation of the tips before it reaches the body temperature inside the GI tract.

We optimized the wax coating conditions for the best wax coverage on the hinges. The distribution of paraffin wax on the wafer is affected by factors such as the wafer's radius, the amount of the viscous liquid, the rotation speed, and rotation time. [3] To achieve optimally uniform wax coverage, we first cut the wafer into quarters to reduce the spinning radius during spin-coating. We spin-coated paraffin wax on microinjectors under different spinning speeds and spinning times, then examined and counted the number of microinjectors with complete wax coverage on the hinges. Table S2 shows the results of the wax deposition optimization experiments. We obtained optimal results where 96% of microinjectors had relatively uniform coverage on their hinges when 500 μL of paraffin wax were deposited at a spin speed of 500 rpm for 10 s and then 1500 rpm for 40 s.

**Table 2. Optimization of wax deposition conditions**

| Wax volume on each quarters 3" wafer | Step 1 | | Step 2 | | The percentage of microinjectors with uniform wax coverage on hinges |
|---|---|---|---|---|---|
| | *Spining speed* | *Time* | *Spining speed* | *Time* | |
| 500 µL | 500 rpm | 10 s | 1000 rpm | 10 s | 12% |
| | | | | 20 s | 13% |
| | | | | 40 s | 24% |
| | | | 1500 rpm | 10 s | 39% |
| | | | | 20 s | 89% |
| | | | | 40 s | 96% |
| | | | 2000 rpm | 10 s | 51% |
| | | | | 20 s | 15% |
| | | | | 40 s | 2% |

**Supporting Note S3: Estimation of the force and pressure exerted by the microinjector tips.**

We estimated the force generated by the actuation of the microinjector tips using maximal force values prior studies that measured the force during the folding of differentially stressed arms using a force-sensing platform [4]. In that study, a 75 nm Cr/ 115 nm Au stress layer on a 100 µm wide hinge produced a force of 4.7 ± 0.9 µN. Our microinjectors have 60 nm Cr/ 100 nm Au and 100 µm hinges, which should generate a similar force value of around 5 µN.

We used the Hertz contact mechanics model to estimate the pressure exerted by the microinjector microtip on the tissue.[5] We approximated the tip of the microinjector as a sphere and the tissue as an elastic half-space. The diameter of the tips was measured from the scanning electron microscope (SEM) images. The microinjectors without a chitosan drug layer have tips of diameter 2R ≈ 3.5 to 5.0 µm. The microinjectors with the chitosan drug layer have tips of diameter 2R ≈ 5.0 to 7.2 µm. We used the value of the maximum force (F) exerted by the microinjector tip as 5 µN according to the above-mentioned literature. From the Hertz model, for the contact between a sphere and a half-space, we obtained the maximum pressure applied on the GI mucosa by the injection tip to be, $P_{max} = \frac{1}{\pi}(\frac{6FE^2}{R^2})^{\frac{1}{3}}$ , where $\frac{1}{E} = \frac{(1-\nu_{tissue}^2)}{E_{tissue}} + \frac{(1-\nu_{tip}^2)}{E_{tip}}$ was calculated using the Youngs' modulus ($E_{tissue}$ = 0.7 MPa, $E_{tip}$ = 55 GPa) and

Poisson's ratio ($v_{tissue} \approx 0.4$, $v_{tissue} = 0.42$) of the GI mucosa[6], and the Au injection tip of the microinjector. Plugging in different injection tip diameters, we found that tips with a chitosan drug patch can apply a pressure of around 0.4 - 0.5 MPa, and the injection tips without a chitosan patch exert a pressure of 0.5 - 0.6 MPa when actuating.

**Supporting Note S4: Validation of the enzyme-linked immunoassay (ELISA) assay.**

We conducted initial pilot experiments to assess the cross-reactivity of the ELISA assay used for the intrinsic rat insulin and to determine the estimate of the amount of human insulin to be delivered to the rats. We found that the ELISA assay does not react to the intrinsic rat insulin, and a substantial amount of insulin can be detected in the rat plasma when 100 mIU of insulin was injected via the intravenous route (Figure S7). We used a dose of 60 mIU of insulin in the delivery experiments with microinjectors for an optimal number of 200 microinjectors in each animal.

**Supporting Note S5: Insulin delivery efficiency of the microinjectors as compared to other published methods**

We examined the insulin delivery efficiency of our microinjector and other GI tract-based insulin delivery mechanisms by comparing the maximum insulin plasma concentration and the insulin dosage per body surface area (BSA) of the testing animals. We extracted the maximum plasma concentrations and initial dosages from the literature. Then we unified their units to *pM* for the plasma concentration and *mg* for the initial dosage. To calculate the BSA of animals, we assume all rats weigh 0.3 kg (the rats in the literature weigh in the range of 0.25 - 0.3 kg), and all pigs weigh 50 kg (the pigs in the literature weigh in the range of 35 - 65 kg). We calculate the BSA of rats[7] by $BSA = \frac{7.47 \times BW^{\frac{2}{3}}}{100}$, and the BSA of pigs[8] by $BSA = \frac{7.98 \times BW^{\frac{2}{3}}}{100}$, BW represents the body weight of the animal in a unit of kg.

#### Supporting Figures

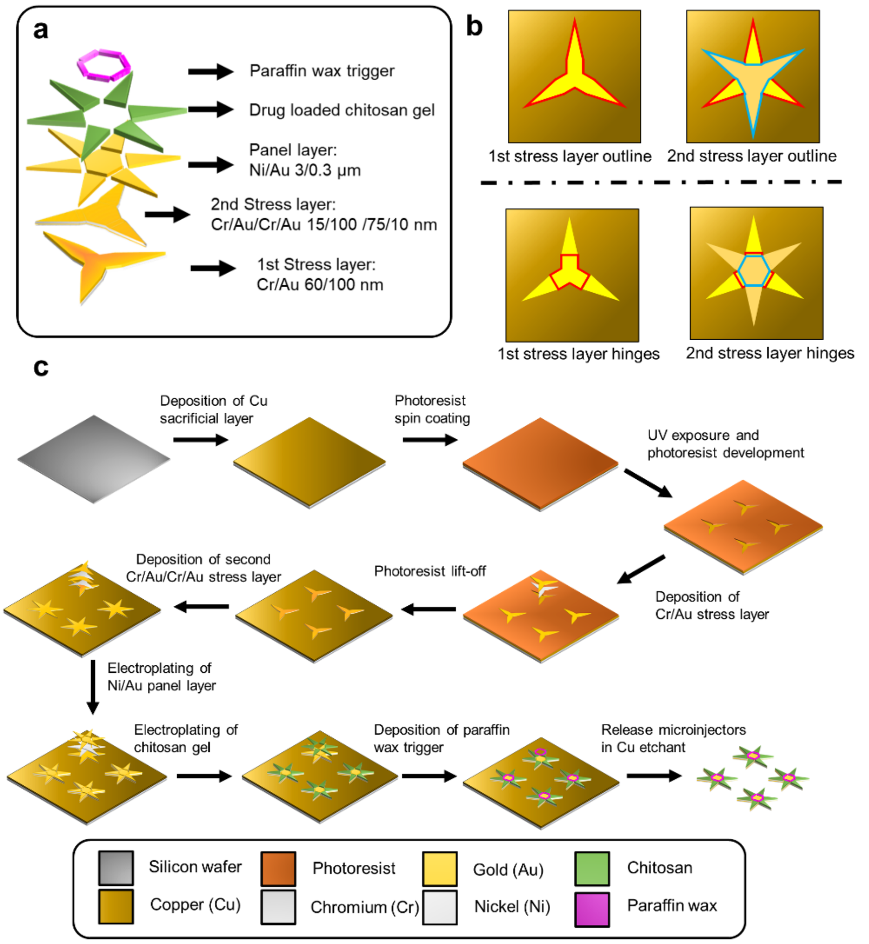


**Figure S1: Schematics showing the design and fabrication of the bidirectional foldable microinjectors. (a)** Schematic showing the magnified view of microinjector's five components. From bottom to top, the components are the 1st stress layer, 2nd stress layer, panel layer, drug loaded chitosan gel and paraffin wax trigger layer. **(b)** Schematic showing the design of the two stress multilayers. The left panel highlights the outline shape of the differentially stressed mutlilayers, and the right panel highlights the shape of the connection part of the differentially stressed multilayers. **(c)** Schematic of step-by-step fabrication of bidirectional foldable microinjectors. The spin coating and UV exposure process is only shown for the first step and this process is repeated for other patterning steps.

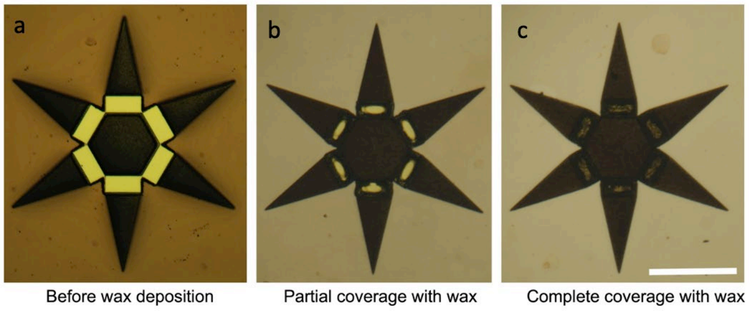


**Figure S2: Optimization of the paraffin wax deposition conditions.** Bright-field optical microscope images showing **(a)** the photoresist mold created by lithography on the hinges of the microinjectors, **(b)** partial deposition of paraffin wax due to non-ideal spin coating conditions, and **(c)** complete coverage of the hinge area with paraffin wax using optimized spin coating conditions. The scale bar is 0.5 mm.

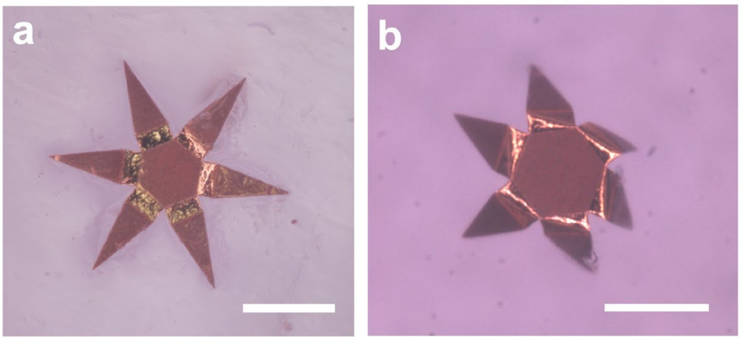


**Figure S3: Microinjectors on tissue-mimicking gelatin.** Bright-field optical microscope images showing a microinjector (a) before actuation and (b) after actuation on 1 kPa stiffness gelatin gel. Note one-sided microinjectors are shown in the images, where all the injector arms actuate in the same direction. The scale bars are 0.5 mm.

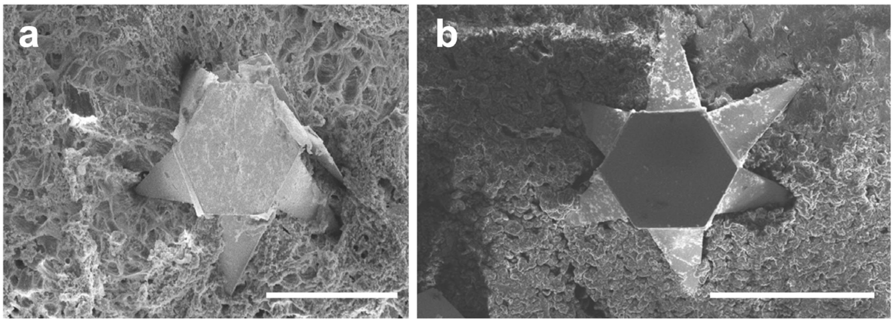


**Figure S4: Microinjectors' penetration of the pig stomach and the colon mucosa.** Scanning electron microscope (SEM) images showing results of *ex vivo* experiments in which one-sided microinjectors can successfully penetrate (a) the pig stomach and (b) the pig colon mucosa. The scale bars are 0.5 mm.

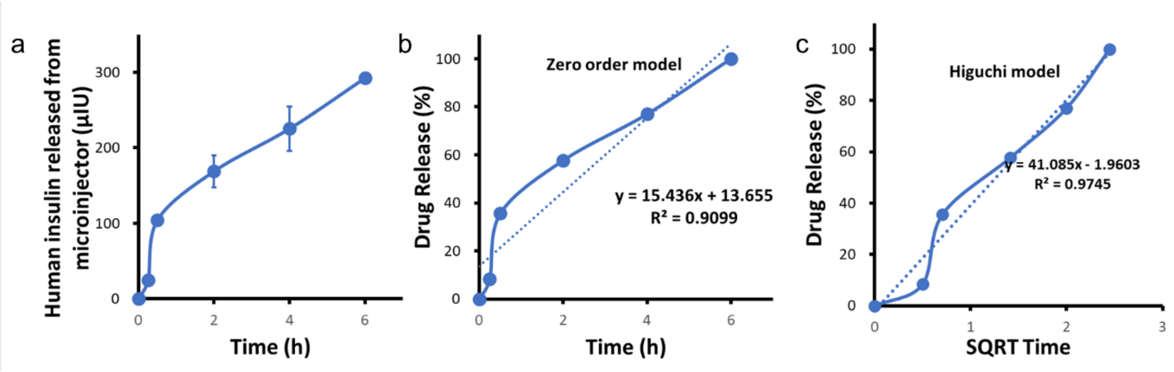


**Figure S5: Microinjector *in vitro* drug release profile.** (a) Plot showing the cumulative measured amount of human insulin released in saline, normalized by the number of injectors. On average, each microinjector can accommodate around 300 micro IU of human insulin. The *in vitro* release experiments were conducted in an oven, set at 37 °C, and repeated four times to generate the mean and standard error of the mean. Plots showing the measured *in vitro* human insulin release profile fit to (b) zero-order and (c) Higuchi model with $R^2$ value of 0.9035 and 0.9745.

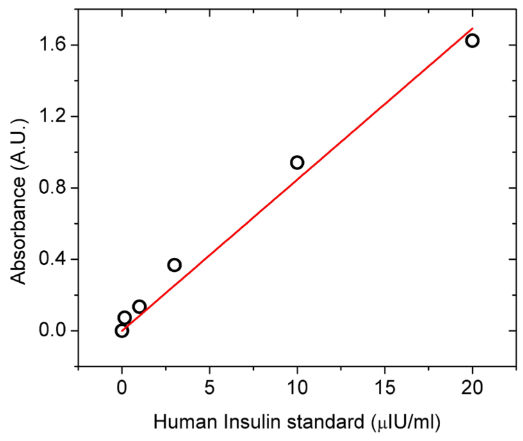


**Figure S6: Standard curve showing absorbance vs. insulin concentration at 450 nm.** A representative standard curve was obtained using a UV-vis spectrophotometer, showing the linear variation of absorbance at 450 nm as a function of the concentrations of the standard insulin solutions used. The red fit line has an $R^2 = 0.9823$, indicating a linear response. The standard solutions were obtained from the manufacturer-supplied human insulin ELISA kit.

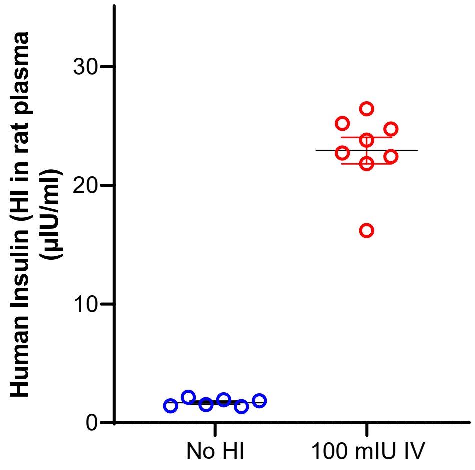


**Figure S7: Validation of the non-specificity of the ELISA assay to intrinsic rat insulin.** A plot of the measured human insulin concentrations in rat plasma for two conditions: no human insulin administered and100 mIU human insulin administered intravenously (IV). We collected the blood plasma 30 minutes after the administration of insulin. Points in the plot represent 1 to 3 repeats from the same animal. N = 2 to 4 animals.

#### Supporting Movie

**Movie SM1:** Video showing thermo-responsive actuation of bidirectional foldable microinjectors sped up 50 times.
